# Growth retardation in a mouse model of Kabuki syndrome 2 bears mechanistic similarities to Kabuki syndrome 1

**DOI:** 10.1101/2023.10.15.562327

**Authors:** Christine W Gao, Wan-Ying Lin, Ryan C Riddle, Sheetal Chopra, Leandros Boukas, Kasper D Hansen, Hans T Björnsson, Jill A Fahrner

**Affiliations:** Department of Genetic Medicine, Johns Hopkins University School of Medicine, Baltimore, MD; Department of Molecular Biology and Genetics, Johns Hopkins University School of Medicine, Baltimore, MD; Department of Orthopaedic Surgery, Johns Hopkins University School of Medicine, Baltimore, MD; Department of Orthopaedics, University of Maryland School of Medicine, Baltimore, MD; Research and Development Service, Baltimore Veterans Administration Medical Center, Baltimore, MD; Department of Biostatistics, Johns Hopkins University School of Public Health, Baltimore, MD; Department of Pediatrics, Johns Hopkins University School of Medicine, Baltimore, MD; Faculty of Medicine, University of Iceland, Reykjavík, Iceland; Landspítali University Hospital, Reykjavík, Iceland

## Abstract

Growth retardation is a characteristic feature of both Kabuki syndrome 1 (KS1) and Kabuki syndrome 2 (KS2), Mendelian disorders of the epigenetic machinery with similar phenotypes but distinct genetic etiologies. We previously described skeletal growth retardation in a mouse model of KS1 and further established that a *Kmt2d^−/−^* chondrocyte model of KS1 exhibits precocious differentiation. Here we characterized growth retardation in a mouse model of KS2, *Kdm6a^tm1d/+^*. We show that *Kdm6a^tm1d/+^* mice have decreased femur and tibia length compared to controls and exhibit abnormalities in cortical and trabecular bone structure. *Kdm6a^tm1d/+^* growth plates are also shorter, due to decreases in hypertrophic chondrocyte size and hypertrophic zone height. Given these disturbances in the growth plate, we generated *Kdm6a^−/−^* chondrogenic cell lines. Similar to our prior *in vitro* model of KS1, we found that *Kdm6a^−/−^* cells undergo premature, enhanced differentiation towards chondrocytes compared to *Kdm6a^+/+^* controls. RNA-seq showed that *Kdm6a^−/−^* cells have a distinct transcriptomic profile that indicates dysregulation of cartilage development. Finally, we performed RNA-seq simultaneously on *Kmt2d^−/−^*, *Kdm6a^−/−^*, and control lines at Days 7 and 14 of differentiation. This revealed surprising resemblance in gene expression between *Kmt2d^−/−^* and *Kdm6a^−/−^* at both time points and indicates that the similarity in phenotype between KS1 and KS2 also exists at the transcriptional level.

## INTRODUCTION

Kabuki syndromes 1 and 2 (respectively, KS1 and KS2; MIM 147920 and 300867) are Mendelian disorders of the epigenetic machinery (MDEMs) (1) with shared features of postnatal growth retardation including short stature and microcephaly, intellectual disability/developmental delay, hypotonia, immune dysfunction, and strikingly similar facial features (2). Patients with either type of Kabuki syndrome are recognizable by their highly arched eyebrows (notched in some cases), elongated palpebral fissures with eversion of the lateral third of the lower eyelid, shortened columella, prominent ears, and persistent fingertip pads. Despite the clinical similarities between KS1 and KS2, their genetic etiologies are distinct.

KS1 accounts for up to 75% of Kabuki syndrome cases (3), and originates from heterozygous, typically *de novo*, pathogenic variants in *KMT2D* (4), which encodes a histone lysine methyltransferase that mono-, di-, and tri-methylates H3K4 (5,6). KMT2D acts at both enhancers and promoters to regulate gene expression. In general, H3K4me1 placed at enhancers and H3K4me3 at promoters are observed at transcriptionally active regions (7). KS2 is comparatively rare, representing 5-8% of Kabuki syndrome (8). Its causative gene, *KDM6A* (9,10), encodes a H3K27 demethylase that preferentially acts upon H3K27me3 and H3K27me2 (11,12). Both modifications are associated with silenced regions of the genome, with H3K27me3 marking transcriptionally repressed promoters, and H3K27me2 coating non-transcribed, intergenic swaths (13,14). *KDM6A* is on the X chromosome, although it is known to escape X inactivation (15). A *KDM6A* homologue, *UTY*, exists on the Y chromosome, but is catalytically inactive (12,16). This imbalance in gene dosage is thus hypothesized to be responsible for the more severe phenotype seen in males with KS2 as opposed to females.

Given the distinct origins of KS1 and KS2, this pair of syndromes provides an interesting study for how divergent epigenetic mechanisms may converge upon similar phenotypes. Fahrner and Björnsson previously proposed the ‘Balance Hypothesis’, such that writers and erasers of epigenetic marks exist in a balance to maintain a normal chromatin state (1). Losing a writer of an activating mark (KMT2D; H3K4 methylation) or losing the eraser of a silencing mark (KDM6A; H3K27 methylation) are both predicted to lead to a closed chromatin state and corresponding transcriptional repression at target genes. Interestingly, chromatin accessibility profiling via ATAC-seq of B cells derived from KS1 and KS2 mouse models showed the converse effect of increased accessibility for both disorders specifically at promoters; such bias was not seen in distal regulatory elements (17). This raised the possibility of indirect effects dominating at the chromatin level – for example, loss of KMT2D or KDM6A may lead to transcriptional silencing of repressors of chromatin accessibility, leading to a paradoxical gain in accessibility. At the transcriptional level, B cells from *Kmt2d^βgeo/+^* and *Kdm6a^tm1d/+^* mice bear considerable overlap in differentially expressed genes when compared to wild-type littermates, suggesting that the same pathways are perturbed in the immune cells of KS1 and KS2. To our knowledge, comparisons between the two disorders have not been made in other affected tissue types.

Growth retardation is a central feature of both KS1 and KS2. The heights of children with KS1 average ≥2 standard deviations below the age-adjusted mean (18). KS2-specific growth curves are unavailable due to the rarity of the disorder, but growth retardation is known to be global and proportionate, with some children also afflicted with microcephaly (3,19,20). We previously used *Kmt2d^βgeo/+^* mice as a model for growth abnormalities in KS1, and showed that disrupted endochondral ossification due to precocious differentiation of chondrocytes contributes towards growth retardation in KS1 (21). *Kmt2d^βgeo/+^* mice had shortened femurs and tibias, and abnormally increased tibial growth plate heights. Cartilaginous growth plates, which expand upon proliferation and hypertrophy of chondrocytes, serve as scaffolds for bone matrix deposition by invading osteoblasts, and are critical to longitudinal bone growth. The finding in *Kmt2d^βgeo/+^* mice of shortened bones in the setting of taller growth plates thus appeared counterintuitive. However, we also saw paradoxically accelerated differentiation and altered transcriptional profile of *Kmt2d^−/−^* chondrocytes *in vitro*. Our data therefore supports the idea that growth retardation in KS1 stems from perturbations to chondrocytes in the growth plates.

Here, we expand our focus to growth retardation in KS2. We leverage the *Kdm6a^tm1d/+^* mouse model of KS2 to characterize skeletal and growth plate abnormalities in this related disorder. In light of our previous findings in KS1, we create *Kdm6a^−/−^* chondrogenic cell lines to investigate chondrocyte differentiation in KS2. We perform RNA-seq of both *Kdm6a^−/−^* and *Kmt2d^−/−^* cells to compare gene expression at different time points and across the two disorders. Our work supports a convergent pattern of gene expression and a common mechanism involving precocious chondrocyte differentiation that leads to growth retardation in both KS1 and KS2.

## RESULTS

### *Kdm6a^tm1d/+^* mice exhibit growth retardation

We utilized a previously described mouse model of KS2, *Kdm6a^tm1d(EUCOMM)Wtsi^* (17), generated using the ‘knockout-first’ targeted construct created by the International Knockout Mouse Consortium (IKMC) (22) and referred to as *Kdm6a^tm1d/+^* henceforth. The *tm1d* allele results from Cre-mediated excision of *Kdm6a* exon 3, which leads to a frameshift and premature termination codon (PTC) in exon 6 of 29 (Figure 1A). Female heterozygous *Kdm6a^tm1d/+^* mice were viable and born at term, but had significantly decreased body weight and length at 8 weeks compared to female *Kdm6a^+/+^* littermates (p=0.007 and p=0.001, respectively) (Figure 1, B and C). Male pups hemizygous for *Kdm6a* were previously described to succumb perinatally (23), and we likewise observed perinatal lethality in *Kdm6a^tm1d/y^* pups. For subsequent experiments, therefore, we focused on heterozygous female mice.

**Figure 1:**
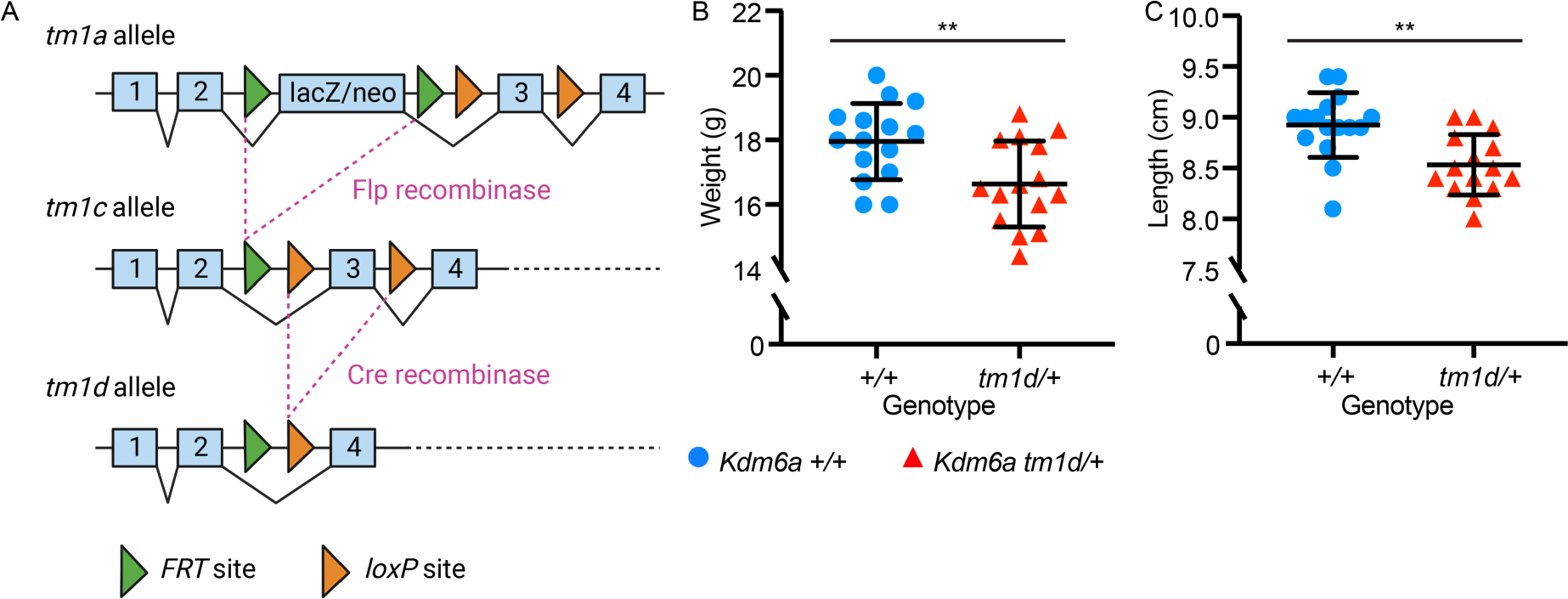
*Kdm6a^tm1d/+^* mice exhibit generalized growth retardation. (A) Schematic of the targeted *Kdm6a* allele as designed by the International Knockout Mouse Consortium, showing *FRT* sites as green arrows and *loxP* sites as orange arrows. Generation of the *tm1d* knockout allele requires excision of exon 3. Compared to *Kdm6a^+/+^* littermates, *Kdm6a^tm1d/+^* mice have significantly decreased (B) body weight (*Kdm6a^+/+^* n=15; *Kdm6a^tm1d/+^* n=15) and (C) body length (*Kdm6a^+/+^* n=16; *Kdm6a^tm1d/+^* n=15). Measurements obtained at 8 weeks of age. Blue circles: *Kdm6a^+/+^*, red triangles: *Kdm6a^tm1d/+^*. **p < 0.01, two-tailed unpaired Student’s t-test. All error bars represent mean ± 1 SD.

To assess growth abnormalities in *Kdm6a^tm1d/+^* mice, we first performed skeletal profiling. *Kdm6a^tm1d/+^* mice had shorter femurs and tibias than *Kdm6a^+/+^* littermates (p=0.0003 and p=0.001, respectively) at 8 weeks of age (Figure 2, A and B). Micro-computed tomography (micro-CT) performed concurrently revealed long bone structural differences in both cortical and trabecular parameters (Figure 2, C–E). *Kdm6a^tm1d/+^* mice had decreased tissue area in the femoral mid-diaphysis (p=0.005) (Figure 2D and F), which was significant even after normalization for femur length (p=0.033) (Supplemental Figure 1). Despite the decreased tissue area, cortical thickness was unaffected in *Kdm6a^tm1d/+^* mice (Figure 2G), leading to a trend towards a higher bone area-to-tissue area percentage (p=0.081) (Figure 2H). Trabecular volume was increased in *Kdm6a^tm1d/+^* mice (p=0.030) (Figure 2, E and I), secondary to a trend towards higher trabecular number (p=0.097) (Figure 2, E and J) and strikingly increased trabecular thickness in *Kdm6a^tm1d/+^* mice (p=0.003) (Figure 2, E and K). These changes suggest disturbances to long bone development due to the heterozygous loss of *Kdm6a*.

**Figure 2:**
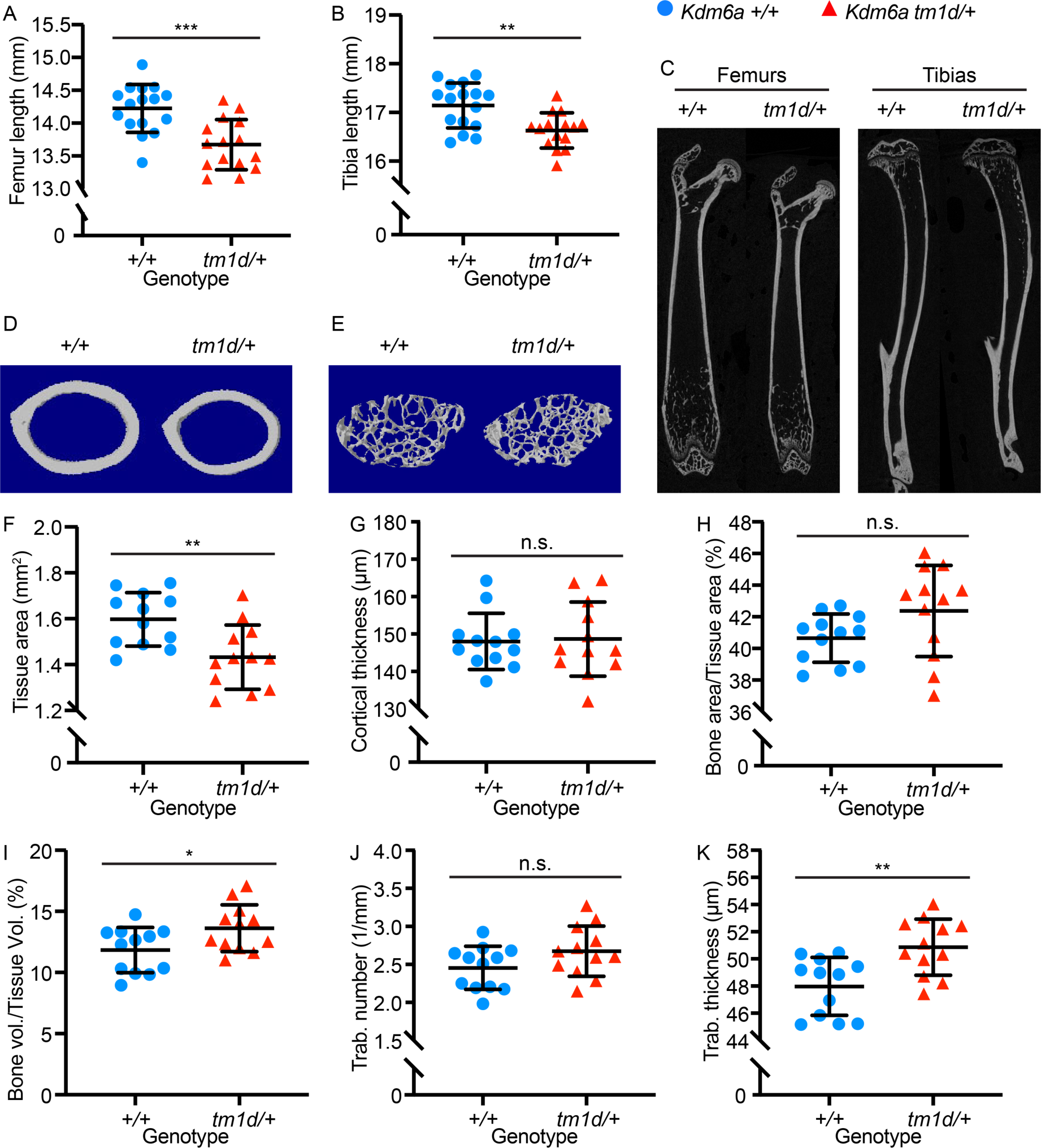
*Kdm6a^tm1d/+^* mice have shortened bones and altered skeletal parameters. *Kdm6a^tm1d/+^* mice have significantly shorter (A) femurs and (B) tibias compared to *Kdm6a^+/+^* littermates (*Kdm6a^+/+^* n=16; *Kdm6a^tm1d/+^* n=15), which is also shown in (C) representative longitudinal micro-CT images of femurs and tibias from *Kdm6a^tm1d/+^* and *Kdm6a^+/+^*. Representative transverse cross-sectional micro-CT reconstructions are displayed for femoral (D) cortical bone and (E) trabecular bone. *Kdm6a^tm1d/+^* mice have decreased (F) femoral cross-sectional tissue area. However, there is no significant difference in (G) cortical thickness or (H) bone area/tissue area percentage, despite *Kdm6a^tm1d/+^* mice displaying a slight upward trend in the latter. (I) Trabecular volume is increased in *Kdm6a^tm1d/+^* mice. While (J) trabecular number does not differ significantly, (K) trabecular thickness is increased in *Kdm6a^tm1d/+^* mice. For (F) through (K), *Kdm6a^+/+^* n=12; *Kdm6a^tm1d/+^* n=12. Blue circles: *Kdm6a^+/+^*, red triangles: *Kdm6a^tm1d/+^*. *p < 0.05, **p < 0.01, ***p < 0.001, two-tailed unpaired Student’s t-test. All error bars represent mean ± 1 SD. n.s., non-significant.

### Growth plate structure is disrupted in *Kdm6a^tm1d/+^* mice

We examined whether perturbed bone development may stem from defects in the growth plate by performing hematoxylin and eosin (H&E) staining of proximal tibial growth plates sectioned in the longitudinal plane at 8 weeks of age (Figure 3A). *Kdm6a^tm1d/+^* mice had an overall decrease in growth plate height (p=0.042) (Figure 3A and B). This was caused by a significant shortening of the hypertrophic zone in *Kdm6a^tm1d/+^* mice compared to wild-type (p=0.0008), although the proliferative zone did not differ from wild-type littermates (Figure 3A and C). Cell counts in both the proliferative and hypertrophic zones were unchanged between the genotypes (Figure 3D). Instead, hypertrophic chondrocyte cell size, as measured by cross-sectional cell area in the longitudinal plane, was

**Figure 3:**
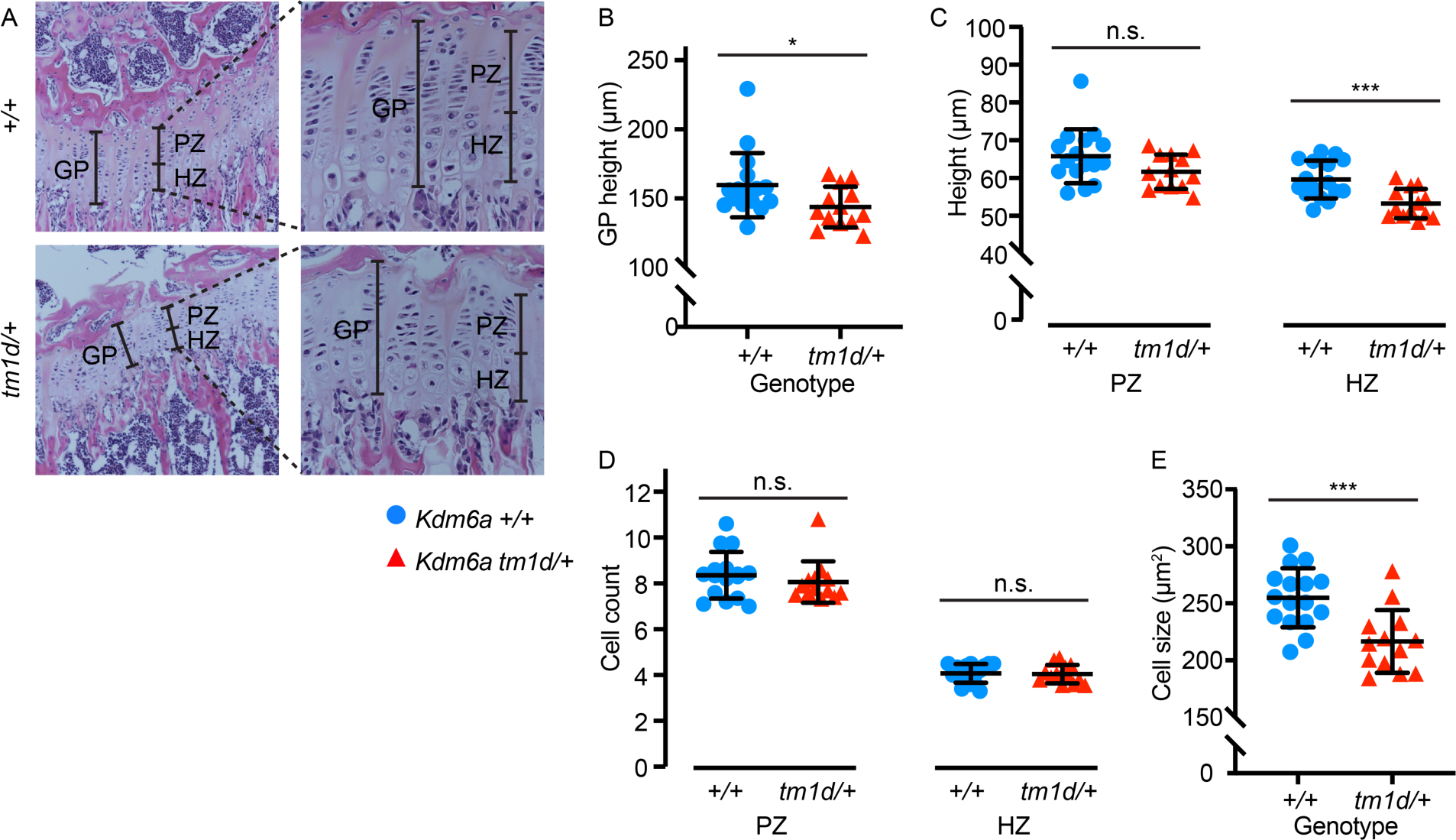
Growth plates have altered structure in *Kdm6a^tm1d/+^* mice. (A) Hematoxylin and eosin staining of *Kdm6a^tm1d/+^* and *Kdm6a^+/+^* tibial growth plates. Left panels: 10x magnification. Insets: 40x magnification. (B) Growth plate height is decreased in *Kdm6a^tm1d/+^* mice. This can be attributed to (C) shorter hypertrophic zones (right panel) in *Kdm6a^tm1d/+^* mice, although proliferative zone heights do not differ (left panel). (D) Cell counts do not differ in either the proliferative (left) or hypertrophic (right) zones, but (E) cross-sectional cell size in the hypertrophic zone is smaller in *Kdm6a^tm1d/+^* growth plates. *Kdm6a^+/+^* n=16; *Kdm6a^tm1d/+^* n=13. Blue circles: *Kdm6a^+/+^*, red triangles: *Kdm6a^tm1d/+^*. *p < 0.05, ***p < 0.001, two-tailed unpaired Student’s t-test. All error bars represent mean ± 1 SD. n.s., non-significant. GP, growth plate. PZ, proliferative zone. HZ, hypertrophic zone. decreased in *Kdm6a^tm1d/+^* mice (p=0.0006) (Figure 3E). Chondrocyte hypertrophy is the main driver of longitudinal bone growth (24). These changes in the growth plate, particularly in the hypertrophic zone, suggest that an abnormality of chondrocyte function is involved in the pathogenesis of growth retardation in KS2.

### *Kdm6a^−/−^* chondrogenic cell lines exhibit precocious differentiation at both phenotypic and transcriptomic levels

To create a tractable *in vitro* model for chondrocyte differentiation and development in KS2, we chose to use ATDC5, a well-characterized female murine teratocarcinoma cell line that is known to undergo chondrogenesis when induced to differentiate (25–27). CRISPR-Cas9 editing was used to target *Kdm6a* exon 6, introducing indels by non-homologous end joining (NHEJ) (Figure 4A). We isolated clones that remained unedited (*Kdm6a^+/+^*) as controls, alongside clones bearing compound heterozygous frameshift mutations (*Kdm6a^−/−^*). All such frameshift mutations led to premature termination codons (PTCs) upstream of the catalytic Jumonji C (JmjC) domain (Supplemental Table 1). For each clone, we excluded off-target edits by Sanger sequencing at the top 3 predicted off-target sites, as identified by partial complementarity to the gRNA (Supplemental Figure 2, A–C). *Kdm6a^−/−^* lines do not express any KDM6A at the protein level, in contrast to *Kdm6a^+/+^* controls (Figure 4B, Supplemental Figure 3A). Intriguingly, *Kdm6a^−/−^* cells did not exhibit higher global levels of H3K27me3 compared to *Kdm6a^+/+^* (Figure 4B, Supplemental Figure 3B). We next differentiated the clones to chondrocytes over the course of 28 days and found that *Kdm6a^−/−^* clones displayed enhanced chondrogenesis. This was visualized by increased Alcian blue staining, which labels glycosaminoglycans (GAGs) secreted by differentiated chondrocytes (Figure 4C) and was subsequently quantified by increased absorbance at 605 nm (Figure 4D). Although we had observed a decrease in cross-sectional cell size in *Kdm6a^tm1d/+^* growth plates (Figure 3E), it was not possible to perform such measurements *in vitro* due to technical limitations imposed by excessive matrix secretion.

**Figure 4:**
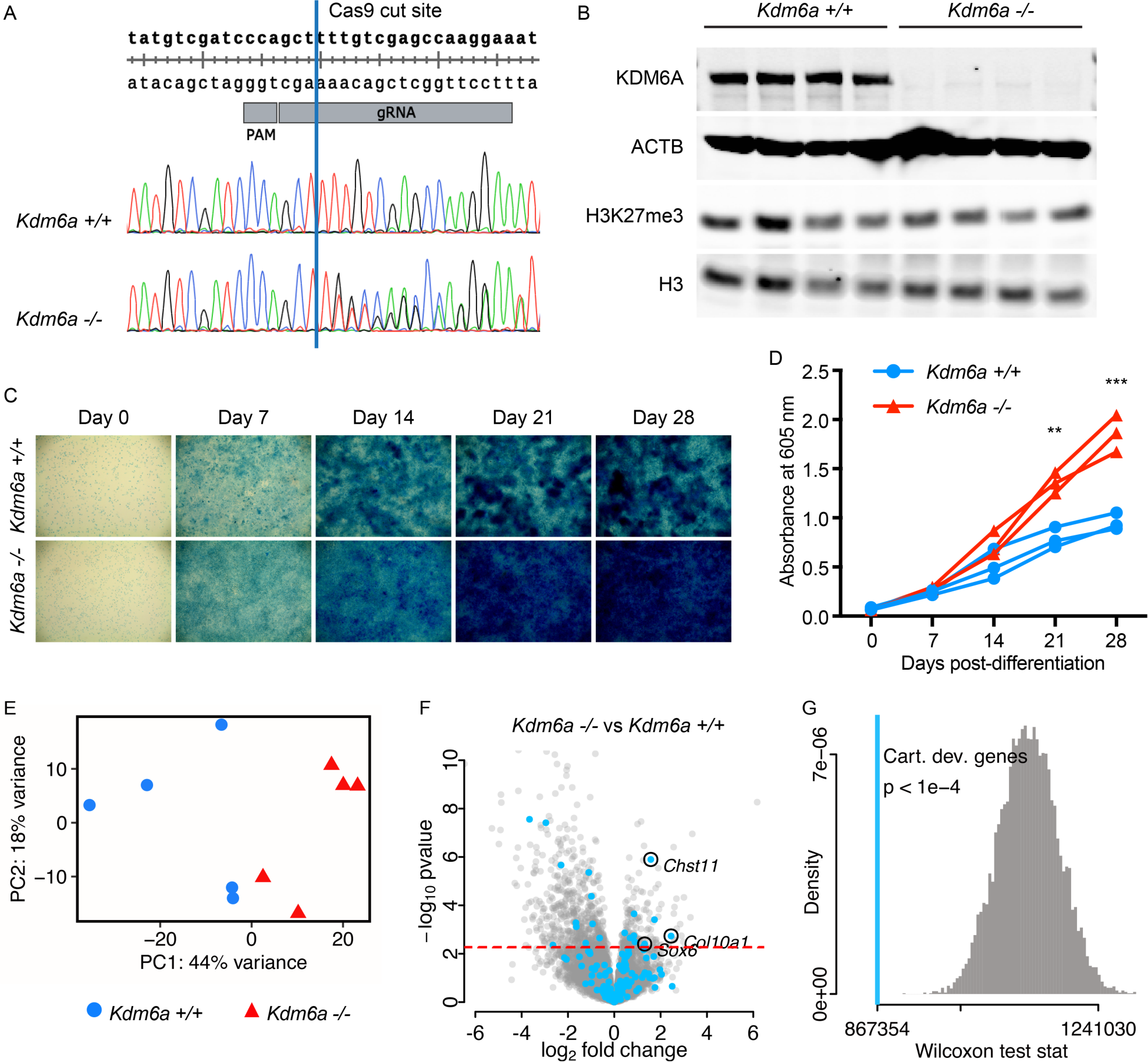
*Kdm6a^−/−^* cells exhibit dysregulated chondrogenesis. (A) CRISPR-Cas9 gRNA target site for generating *Kdm6a^−/−^* cell lines. Representative chromatograms from an unedited *Kdm6a^+/+^* clone and a compound heterozygous *Kdm6a^−/−^* clone, which shows double peaks following the intended Cas9 cut site (blue line). (B) *Kdm6a^−/−^* cells do not express KDM6A protein, and H3K27me3 levels are unaltered from *Kdm6a^+/+^* control lines. (C) Representative Alcian blue staining images (4x magnification) from a 28-day time course of *Kdm6a^−/−^* and *Kdm6a^+/+^* cell lines. (D) *Kdm6a^−/−^* cells lines have increased uptake of Alcian blue stain. Each line represents a single technical replicate, comprising the average of 3-4 biological replicates. (E) Principal component analysis of *Kdm6a^−/−^* and *Kdm6a^+/+^* RNA-seq samples collected at Day 14 of differentiation (*Kdm6a^+/+^* n=5; *Kdm6a^−/−^* n=5). Blue circles: *Kdm6a^+/+^*, red triangles: *Kdm6a^−/−^*. (F) Volcano plot for *Kdm6a^−/−^* versus *Kdm6a^+/+^* samples, with MGI cartilage development genes highlighted in light blue. False discovery rate (FDR) = 0.1 (red dashed line). (G) Wilcoxon rank sum test statistic for MGI cartilage development genes (light blue line, p < 1e-4) and simulated distribution of test statistics (gray). **p < 0.01, ***p < 0.001, two-tailed unpaired Student’s t-test. Cart. dev. genes, cartilage development genes.

To identify the transcriptional changes behind the chondrogenic phenotype in *Kdm6a^−/−^* cells, we performed RNA-seq on 5 *Kdm6a^+/+^* and 5 *Kdm6a^−/−^* clones at day 14 of chondrocyte differentiation. We chose this time point immediately prior to the increase in Alcian blue uptake in *Kdm6a^−/−^* because changes in gene expression are expected to precede phenotypic manifestation. Principal component analysis showed that the clones clustered according to genotype, with principal component 1 (PC1) accounting for 44% of inter-sample variance (Figure 4E). A total of 736 genes were differentially expressed at the 10% false-discovery rate (FDR) level (Supplemental Appendix 1). Of these, 541 genes were downregulated in *Kdm6a^−/−^* chondrocytes and only 195 genes were upregulated (Figure 4F), suggesting a skew towards repression of gene expression – or possibly, loss of activation.

We also found perturbations in chondrogenic pathways at the transcriptional level. From a list of 200 genes annotated by the Mouse Genome Informatics (MGI) database as involved in cartilage development (see Methods) (Supplemental Appendix 2), 164 genes were expressed in our RNA-seq dataset. These genes tended to have greater absolute fold-change in expression level than non-chondrogenic genes, suggesting that cartilage development could be collectively dysregulated in *Kdm6a^−/−^* cells (Supplemental Figure 4). To quantify this effect, we showed that the p-value rank-sum of this group of genes was significantly different from the distribution of p-value rank-sums obtained from selecting groups of 164 genes at random (p < 1×10^−4^) (Figure 4G). Specific genes of interest include *Col10a1*, which was significantly overexpressed in *Kdm6a^−/−^* cells with a log2(fold-change) of 2.45 (Figure 4F) (28). *Chst11*, a carbohydrate sulfotransferase responsible for the production of chondroitin sulfate, the major proteoglycan component of cartilage (29), was also significantly upregulated. Finally, *Sox6*, a transcription factor known to regulate chondrogenesis by promoting hypertrophy and organization of the growth plate (30), was also overexpressed. These RNA-seq findings support that loss of KDM6A causes transcriptional alterations leading to excessive chondrogenesis.

### *Kmt2d^−/−^* and *Kdm6a^−/−^* cell lines share a transcriptional profile indicative of excessive chondrogenesis

The similarities of precocious differentiation in both KS1 and KS2 chondrocytes prompted a direct comparison of their transcriptional profiles. We had previously generated *Kmt2d^−/−^* and control *Kmt2d^+/+^* clones from the same ATDC5 parental cell line by CRISPR-Cas9 gene editing. RNA-seq at Day 7 of chondrogenic differentiation revealed striking differential gene expression in *Kmt2d^−/−^* lines compared to *Kmt2d^+/+^* (21). To account for the difference in timepoints and minimize batch effects, we concurrently cultured *Kmt2d^−/−^* and *Kdm6a^−/−^* lines and their respective wild-type control lines. Poly-adenylated RNA samples were sequenced from all samples at both Days 7 and 14 of differentiation. We first validated these results with our prior datasets. We stratified genes on the basis of significance at an FDR cutoff of 10% in our previous data, obtained at Day 7 for KS1 (21). In our new dataset, we contrasted Day 7 *Kmt2d^−/−^* samples with Day 7 *Kmt2d^+/+^*, and the new experiment indicates a strong enrichment of genes that were also significant in prior data (Supplemental Figure 5A). Furthermore, the fold-change directionality was concordant for 864 out of 866 genes significant in both datasets (99.8%) (Supplemental Figure 5B). For KS2, we performed a similar analysis stratifying genes based on prior data collected at Day 14, and again noted significant enrichment for the same genes that were previously identified (Supplemental Figure 5C), as well as high concordance in the direction of fold-change (403 out of 404 genes significant in both data sets; 99.8%) (Supplemental Figure 5D). Altogether, this confirmed that our new data aligns with prior findings where applicable, and allowed us to directly compare KS1 and KS2.

To investigate how factors such as genotype and differentiation stage affected sample clustering, we performed principal component analysis (PCA). This indicated separation roughly based on genotype, with principal component 1 (PC1) accounting for 32% of variance (Figure 5A). Interestingly, there was substantial variation among wild-type samples. *Kmt2d^+/+^* lines that were isolated as matched controls for *Kmt2d^−/−^* clones appeared to separate along principal component 2 (PC2, 18% of variance) away from *Kdm6a^+/+^* controls isolated contemporaneously with *Kdm6a^−/−^* lines, despite the wild-type status of both groups. We have previously confirmed by Sanger sequencing that *Kmt2d^+/+^* and *Kdm6a^+/+^* lines do not bear edits in the targeted loci, or in the top predicted CRISPR/Cas9 off-target sites (Supplemental Figure 2, A–C). Furthermore, PC1 accounts for nearly twice the variance as PC2 (32% versus 18%). However, in case of divergence between the two sets of wild-type samples over the course of passaging, we elected to make all further comparisons between knockout lines only with their respective matched wild-type controls, henceforth referred to as *Kmt2d^+/+^* for KS1 and *Kdm6a^+/+^* for KS2.

**Figure 5:**
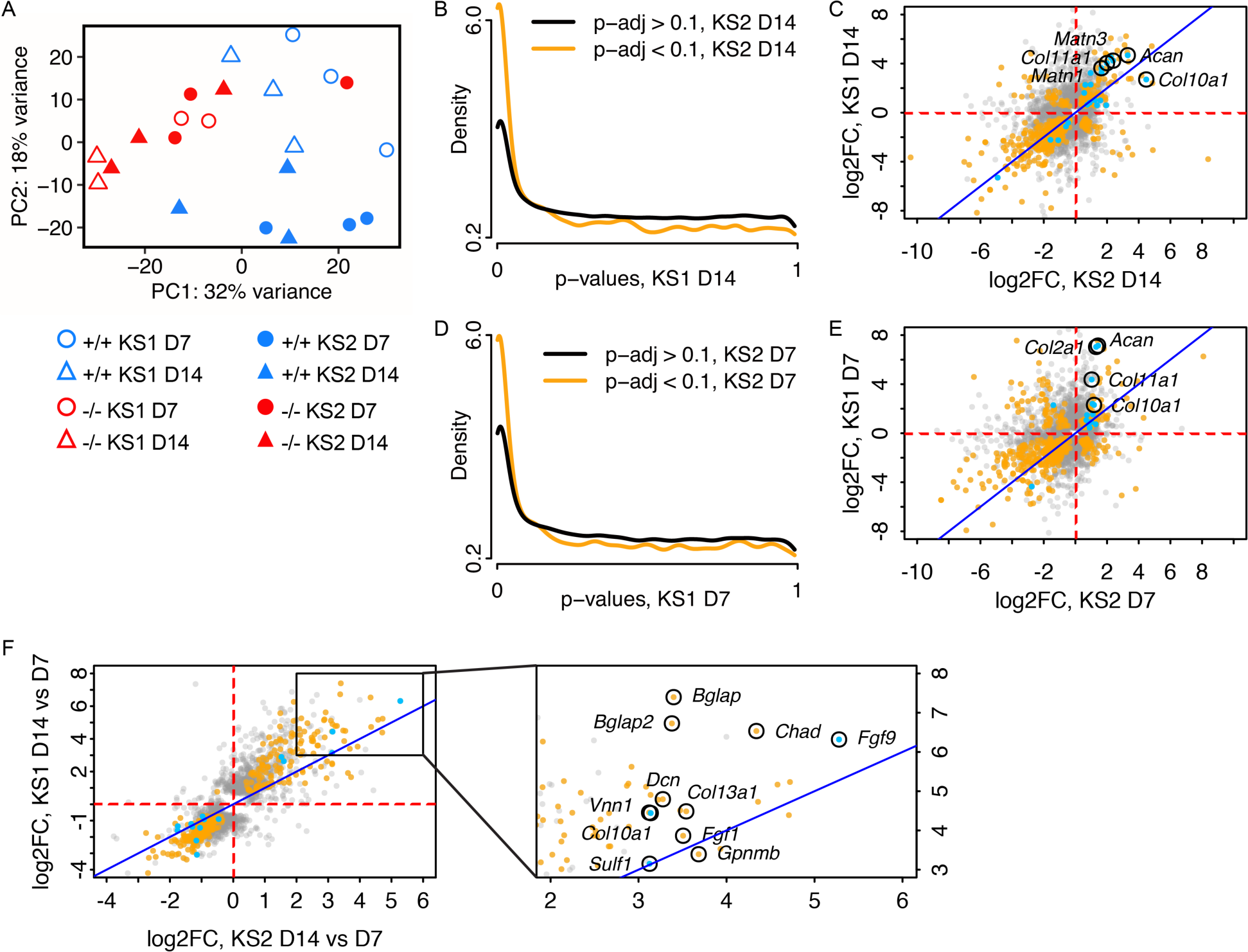
*Kmt2d^−/−^* and *Kdm6a^−/−^* chondrocytes bear similar transcriptomic profiles. (A) Principal component analysis of Day 7 and Day 14 RNA-seq samples from *Kmt2d^−/−^* (KS1; n=2), *Kdm6a^−/−^* (KS2; n=3), and wild-type control cell lines *Kmt2d^+/+^* (KS1; n=3) and *Kdm6a^+/+^* (KS2; n=3), respectively. Blue symbols represent wild-type; red symbols represent knockout lines. Circles represent Day 7; triangles represent Day 14. Open symbols represent KS1 (*Kmt2d^−/−^* and *Kmt2d^+/+^*) and solid symbols represent KS2 (*Kdm6a^−/−^* and *Kdm6a^+/+^*). (B) Conditional p-value histogram displaying p-values from the contrast of *Kmt2d^−/−^* versus *Kmt2d^+/+^* at Day 14 (KS1 D14), stratified by gene-wise significance at Day 14 in the contrast of *Kdm6a^−/−^* versus *Kdm6a^+/+^* (KS2 D14; orange line, p-adj < 0.1) or non-significance (black line, p-adj > 0.1). (C) Scatter plot of gene-wise log2(fold-changes) at Day 14 in the *Kdm6a^−/−^* versus *Kdm6a^+/+^* contrast (KS2) and the *Kmt2d^−/−^* versus *Kmt2d^+/+^* contrast (KS1). Orange: gene significant in both KS1 and KS2; gray: gene is significant in either KS1 or KS2, but not both; blue: gene is annotated in MGI as involved in cartilage development. Identity line (x=y) is displayed in dark blue. The (D) conditional p-value histogram is also displayed for Day 7, as is the (E) scatter plot of gene-wise log2(fold-changes) at Day 7. (F) Scatter plot of gene-wise log2(fold-changes) between Day 14 and Day 7, for KS1 and KS2. Inset panel shows a magnified area of the plot.

Notably, the PCA plot also showed that, for the most part, *Kmt2d^−/−^* samples cluster tightly with *Kdm6a^−/−^* samples regardless of timepoint (Figure 5A). To determine the overlap in gene expression patterns between KS1 and KS2 at Day 14, we compared *Kmt2d^−/−^* and *Kdm6a^−/−^* lines against *Kmt2d^+/+^* and *Kdm6a^+/+^* samples, respectively (differentially expressed genes are listed in Supplemental Appendices 3 and 4). Gene-wise p-values in KS1 were stratified by their significance in KS2 at an FDR cutoff of 10% (Figure 5B). This revealed that genes with low p-values in KS2 are likewise enriched for low p-values in KS1, suggesting a high degree of similarity between the chondrocyte transcriptomes of the two disorders. The fold-change directionality from wild-type was also extremely concordant between KS1 and KS2; of the 578 genes significant in both conditions, 139 genes were upregulated in both KS1 and KS2 and 362 of genes were downregulated, leaving only 77 genes discordant between the two conditions (Figure 5C). Mutually upregulated genes of interest include: *Acan*, which encodes aggrecan, a major proteoglycan component in cartilage; *Matn1* and *Matn3*, both encoding matrilin proteins that are part of the cartilage matrix; and several collagen genes including *Col10a1* and *Col11a1*. We also found a high degree of overlap in differentially expressed genes between KS1 and KS2 at Day 7 (Figure 5D; Supplemental Appendices 5 and 6), although fold change directionality was generally less concordant at Day 7 than at Day 14 (Figure 5E). *Acan, Col10a1, Col11a1,* and *Col2a1,* an early marker for chondrogenesis (31), were significantly upregulated in both conditions at Day 7. These results show that KS1 and KS2 have similar transcriptomic profiles, which support increased chondrocyte differentiation compared to wild-type controls.

We had performed RNA-seq at two time points, on Day 7 and Day 14 of differentiation, in order to examine the progression in gene expression in KS1 and KS2. This allowed us to assess whether the chondrocyte differentiation program differs between KS1 and KS2. We compared *Kmt2d^−/−^* and *Kdm6a^−/−^* samples collected at Day 14 against those collected at Day 7, and there was increased expression of a number of chondrogenic and osteogenic genes at the later time point (Figure 5F). *Fgf9*, a transcription factor that promotes chondrocyte hypertrophy, had 38.8-fold higher expression at Day 14 in KS1 and 79.8-fold increase in KS2, compared to Day 7. *Bglap* and *Bglap2*, which encode calcium-chelating protein hormones produced exclusively in bone tissue, were also mutually upregulated – particularly so in KS1. The overall pattern of gene expression changes from Day 7 to Day 14 indicated an increasingly chondrogenic and osteogenic environment in both *Kmt2d^−/−^* and *Kdm6a^−/−^* cells. Interestingly, the similarity in gene-wise fold-changes between KS1 and KS2 is greater when considering the change between time points rather than within a given time point (Figures 5C, E, and F). This suggests that the chondrogenic differentiation program is largely shared between KS1 and KS2.

## DISCUSSION

Growth abnormalities are a hallmark feature of Mendelian disorders of the epigenetic machinery (MDEMs). Here we characterized growth retardation in a mouse model of KS2. Similar to individuals with KS2, *Kdm6a^tm1d/+^* mice had reduced body length and weight. We also showed that *Kdm6a^tm1d/+^* mice have shorter femurs and tibias alongside other skeletal abnormalities. Unlike in the previously published *Kmt2d^βgeo/+^* model of KS1 (21), *Kdm6a^tm1d/+^* mice display a decrease in growth plate height. At the cellular level, *Kdm6a^−/−^* cell lines show precocious differentiation of chondrocytes, similar to *Kmt2d^−/−^* lines. The increase in chondrogenesis is captured in the transcriptomic profiles of both *Kmt2d^−/−^* and *Kdm6a^−/−^* cells. To our knowledge, this is the first comparison of gene expression between KS1 and KS2 in a cell type related to growth retardation.

We used cell lines that were null for *Kmt2d* or *Kdm6a* because it is generally accepted that complete loss of a protein produces a more profound effect than heterozygosity. This eased initial characterization of the phenotype and gene expression changes. However, most MDEMs – including KS1 – are dominant disorders; KS2 is X-linked dominant. Owing to the critical functions of the epigenetic machinery, constitutional homozygous pathogenic variants have been shown to be lethal in mouse models (6,32), although conditional knockouts may be viable. Therefore, *Kmt2d^−/−^* and *Kdm6a^−/−^* cell lines likely capture the extreme end of the phenotypic and transcriptomic spectrum, which was displayed *in vivo* by our heterozygous mouse models that mimic human disease.

### *Kdm6a^tm1d/+^* mice display a defect in longitudinal and appositional bone growth

Bone growth is closely coupled with mechanical load. Mice with lower body mass and smaller skeletal frames, as we see with the *Kdm6a^tm1d/+^* genotype, thus proportionally have lower femoral cross-sectional tissue area. However, we showed that *Kdm6a^tm1d/+^* mice have a significantly decreased tissue area even after normalization for shorter femur length. As tissue area can only be expanded by osteoblast activity at the periosteal surface, the lack of appropriate tissue area growth suggests that osteoblasts may be another afflicted cell type in KS2. Interestingly, femoral cortical thickness was unchanged in *Kdm6a^tm1d/+^* mice and trabecular volume was increased. Decreased resorptive capability by osteoclasts at the endosteal and trabecular surfaces may also contribute towards the skeletal phenotype in KS2. Although we focus on chondrocytes here, our *in vivo* findings suggest that future studies of other cell types involved in bone growth – specifically osteoblasts and osteoclasts – would be important to expand understanding of growth retardation in KS2.

### Overlap between KS1 and KS2 is apparent in gene expression profiles

An extension of the Balance Hypothesis (1) supposes that impaired function of distinct components of the epigenetic machinery may yet converge upon the same phenotype by achieving a shared chromatin state and gene expression. We found that despite the distinct genetic etiologies of KS1 and KS2, downstream gene expression patterns appear to be shared, which has also been noted by others. Luperchio et al. used RNA-seq on B lymphocytes isolated from mouse models of KS1 and KS2 and found a high degree of overlap in differential gene expression (17). Several groups have developed a genome-wide DNA methylation signature in peripheral blood that can differentiate individuals with KS1 from healthy controls (33–35). Interestingly, this signature also identifies KS2 samples due to the high degree of overlap, and has even aided in diagnosis (33,36,37). As gene expression is dependent upon DNA methylation status, this suggests that there is a shared ‘Kabuki transcriptome’ in peripheral blood. Relative to wild-type controls, we showed striking concordance in gene expression between *Kmt2d^−/−^* and *Kdm6a^−/−^* cells at multiple stages of chondrocyte differentiation. The transcriptional signatures of these genotypes were both suggestive of precocious chondrogenesis. This also matched our cellular phenotype of increased Alcian blue staining in *Kdm6a^−/−^* lines, described here, and our previously published findings in *Kmt2d^−/−^* lines (21).

### KS1 and KS2 may lead to growth retardation through distinct pathways

Several pieces of data indicate that the mechanism of growth retardation in KS1 may still differ from KS2 in subtle ways. Our prior work identified *Shox2* as a direct target of Kmt2d (21). *Shox2* is expressed in mouse growth plates, and conditional loss of *Shox2* in chondrocytes causes shortened limbs (38,39). *Kmt2d^−/−^* cells exhibited depletion of H3K4me3 at the *Shox2* promoter relative to *Kmt2d^+/+^* lines, concurrent with a 4-fold downregulation of *Shox2* expression. The decrease in SHOX2 led to a disinhibition of *Sox9* expression in *Kmt2d^−/−^* cells. As SOX9 is a pro-chondrogenic transcription factor (40,41), we proposed that this pathway is implicated in the precocious differentiation of chondrocytes, which we have shown to be a common feature of KS1 and KS2. Yet in *Kdm6a^−/−^* cells, neither *Shox2* nor *Sox9* is differentially expressed relative to *Kdm6a^+/+^* controls, at either Day 7 or Day 14 of chondrogenic differentiation. This suggests that the *Shox2-Sox9* pathway is not a driving factor in precocious chondrogenesis in KS2. Notably, even in *Kmt2d^−/−^* cells, repletion of *Shox2* by lentiviral overexpression was insufficient to restore Alcian blue staining to wild-type levels (21), indicating that additional pathways – possibly shared between KS1 and KS2 – contribute towards this phenotype.

A caveat to these conclusions is that our *Kmt2d^+/+^* and *Kdm6a^+/+^* control lines, both previously confirmed to be wild-type, appear to differ at baseline. Whereas this may be due to passaging effects, as the clones were derived on separate instances, it remains possible that the aforementioned gene expression differences between KS1 and KS2 could be an artifact of our imperfect cell models. However, detailed examination of skeletal profiles in KS1 and KS2 mouse models also reveals differences despite the preserved global picture of growth retardation. Compared to wild-type littermates, *Kmt2d^βgeo/+^* and *Kdm6a^tm1d/+^* mice have significantly lower body weight and body length, and shorter femurs and tibias (21). Unlike *Kdm6a^tm1d/+^* mice, however, *Kmt2d^βgeo/+^* mice did not have any significant difference in tissue area, and trabeculae were thinner compared to wild-type. In the tibial growth plate, *Kdm6a^tm1d/+^* mice had reduced hypertrophic chondrocyte size, leading to a significant decrease in hypertrophic zone and overall growth plate height. This was the opposite of growth plate findings in *Kmt2d^βgeo/+^* mice. These data definitively point to chondrocytes as a critical cell type in the pathogenesis of growth retardation for both KS1 and KS2. However, it remains a question how to reconcile opposing growth plate data with the ultimate shared phenotype of shortened femurs and tibias.

### Chondrocyte-to-osteoblast transdifferentiation remains to be explored in KS1 and KS2

Altered cell differentiation is a common disease process in this class of disorders, as described in other MDEMs (21,42,43). Cell type-specific gene expression is generally controlled through epigenetics, and prior studies indicate that disruptions to the epigenetic machinery impact cell fate and commitment (44). The classical model of endochondral ossification describes apoptosis of hypertrophic chondrocytes and subsequent invasion of the residual cartilage template by osteoblasts. Accelerated chondrocyte differentiation, as appears to occur in KS1 and KS2, could therefore impact downstream processes of bone formation. Not only might an altered epigenome hasten or delay differentiation, it is also conceivable that the barriers between cell types may become increasingly fluid. Chondrocytes and osteoblasts were originally believed to be mutually exclusive, committed cell fates resulting from differentiation of mesenchymal stem cells (45). However, lineage tracing experiments using transgenic mice have now led to several hypotheses for chondrocyte-to-osteoblast transdifferentiation (46). Whereas some groups propose transdifferentiation at the immature chondrocyte stage (47), others suggest dedifferentiation of hypertrophic chondrocytes followed by redifferentiation (48–50), and yet additional data may support direct transdifferentiation from hypertrophic chondrocytes into mature osteoblasts (51–53). A disruption in this complex interplay of cell fates could plausibly contribute to growth plate differences in KS1 and KS2. We noted an extreme upregulation of *Bglap* and *Bglap2* between Day 7 and Day 14 of chondrocyte differentiation in *Kmt2d^−/−^* cell lines in particular. Alternatively named *Osteocalcin*, these genes are regarded to be osteoblast-specific (54), suggesting that transdifferentiation may be occurring. *Kdm6a^−/−^* and wild-type cell lines also increased expression of these genes over the course of differentiation, although to a lesser degree. ATDC5 cells are an established *in vitro* model system for chondrogenesis (25–27). However, it would be ideal to further investigate *in vivo* whether KS1 and/or KS2 change the potential for chondrocyte-to-osteoblast transdifferentiation, as an imbalance in these cell types would impact the bone environment and appositional bone growth.

### KMT2D and KDM6A co-exist in the COMPASS complex

The Balance Hypothesis is an appealing explanation for the clinical resemblance of KS1 and KS2, but it is important to consider alternative hypotheses as well. In support for the former, Björnsson et al. have shown that *Kmt2d^βgeo/+^* mice have genome-wide depletion of H3K4me3, which is the expected direct consequence of reduced KMT2D activity (55). This was reversible upon treatment with a histone deacetylase inhibitor (HDACi), AR-42, which also ameliorated neurologic findings of KS1 in the mice. These data, in combination with the Kabuki syndrome DNA methylation signatures (33–35), point towards an epigenetic etiology for these disorders. Luperchio et al. also found that *Kmt2d^βgeo/+^* and *Kdm6a^tm1d/+^* B lymphocytes both tend towards increased promoter accessibility (17). On the one hand, this is supportive of KS1 and KS2 having a shared chromatin state that contributes to overlapping gene expression, which we saw in our RNA-seq data comparing *Kmt2d^−/−^* and *Kdm6a^−/−^* cells. Yet, loss of either a writer of H3K4me3 or an eraser of H3K27me3 would be expected to have the opposite effect on chromatin accessibility. Two potential possibilities (which are not mutually exclusive) include: 1. An indirect effect that is downstream of KMT2D or KDM6A catalytic activity drives the determination of chromatin state, or 2. The role of KMT2D and/or KDM6A in the pathogenesis of Kabuki syndrome stems from a non-catalytic function in one or both of these proteins.

Our western blot results showing that global H3K27me3 levels are unaltered even in *Kdm6a^−/−^* cells with no residual KDM6A protein. This is not entirely unsurprising, as the KDM6 family also includes KDM6B, which shares 70% protein sequence identity with KDM6A and also has H3K27 demethylase activity (12). However, it raises the possibility that the predominant role of KDM6A in KS2 may be non-catalytic. KMT2D and KDM6A coexist in the COMPASS complex, which broadly serves as an antagonist to gene repression by Polycomb group proteins throughout development (56). The reduced dosage of either KMT2D or KDM6A could disrupt stoichiometry or destabilize the COMPASS complex – and perhaps the shared features of KS1 and KS2 are rooted in the destabilization of COMPASS rather than the individual enzymatic functions of KMT2D and KDM6A. Domain-swapping experiments have shown that KDM6A has a non-catalytic capacity to recruit the histone acetyltransferase p300 to co-localize with COMPASS at enhancer elements (57). Likewise, catalytic-dead KDM6A is sufficient to rescue differentiation in *Kdm6a*-knockout mouse embryonic stem cells (23,58), establishing that KDM6A plays important non-enzymatic roles. It would be interesting to consider the formation and genomic localization of the COMPASS complex in the absence or reduced dosage of KMT2D, as well as KDM6A. Should the catalytic role of KMT2D also be implicated in the pathogenesis of KS2, it may unlock the possibility of treatment with pharmacological and/or dietary therapies, which have already been shown to improve neurological outcomes in *Kmt2d^βgeo/+^* mice (55,59,60).

In this work, we aimed to further understand the mechanisms involved in growth retardation for two related MDEMs, KS1 and KS2. We showed that while skeletal and growth plate findings differ in certain aspects between mouse models of KS1 and KS2, chondrocytes bearing *Kmt2d* or *Kdm6a* null mutations exhibit a shared phenotype of precocious differentiation. Importantly, the similarity in transcriptional profiles raises the tantalizing possibility of developing a treatment to target both disorders.

## METHODS

### Animals

The *Kdm6a^tm1a(EUCOMM)Wtsi^* allele (*tm1a*) was designed by the International Knockout Mouse Consortium (IKMC) and consisted of a *FRT*-flanked LacZ-neomycin resistance cassette (*βgeo*), upstream of the *lox*P-flanked *Kdm6a* exon 3. *Kdm6a^tm1a/+^* females were obtained by the Björnsson lab from the European Mutant Mouse Archive (EMMA) and crossed with B6.Cg-Tg(ACTFLPe)9205Dym/J males (Strain No. 005703, The Jackson Laboratory), which express *FLP1* driven by the *Actb* promoter. This excised the *βgeo* cassette and generated the *tm1c* allele. *Kdm6a^tm1c/tm1c^* females were next crossed with B6.C-Tg(CMV-cre)1Cgn/J males (Strain No. 006054, The Jackson Laboratory), which carry a human CMV-driven *cre* transgene integrated on chromosome X. *Cre*-mediated excision of *Kdm6a* exon 3 resulted in the *tm1d* allele. *Kdm6a^tm1d/+^* females were backcrossed to C57BL/6J males (Strain No. 000664, The Jackson Laboratory) to remove *FLP1* and *cre* transgenes and to maintain the strain subsequently. Genotyping was performed by PCR. All mice were in the care of Johns Hopkins Research Animal Resources. Up to 5 mice were housed in each barrier cage with *ad libitum* access to autoclaved feed (Envigo Teklad 2018SX) and reverse osmosis-filtered, hyperchlorinated water. A cotton square nestlet (Envigo Teklad 6060/6105) and autoclaved corncob bedding (Envigo Teklad 7092/7097) were provided for each cage. Cages were changed every 2 weeks under aseptic conditions. Cage racks were ventilated with HEPA-filtered and humidified air. Animal rooms were maintained on standard light/dark cycles at regulated temperatures. Mice were euthanized by halothane inhalation (Sigma-Aldrich B4388), following the AVMA Guidelines for the Euthanasia of Animals, 2020 edition (https://olaw.nih.gov/policies-laws/avma-guidelines-2020.htm).

### High-resolution micro-computed tomography

Mice were euthanized at 8 weeks of age. Femurs and tibias were dissected and fixed with 4% paraformaldehyde in 1x PBS for 48-72 hours at 4°C, then transferred to 70% ethanol. Digital calipers were used to measure bone length. Imaging was performed following the guidelines of the American Society for Bone and Mineral Research (61). A Bruker Skyscan 1172 desktop microcomputed tomography system was used to scan bones at 65 keV and 152 µA with a 0.5 mm aluminum filter at an isotropic voxel size of 10 µm. Image reconstruction was performed with nRecon (Bruker), and analysis was done with CTAn software (Bruker). A 500 µm region of interest (ROI) centered around the femoral mid-diaphysis was assessed for cortical bone measurements. Trabecular measurements were collected 500 µm proximal to the distal femoral growth plate in a 2 mm ROI.

### Growth plate measurements

Femurs and tibias dissected from 8 week-old mice were fixed with 4% paraformaldehyde in 1x PBS for 48-72 hours at 4°C, then decalcified by continual agitation in a 14% EDTA solution, pH 7.4 (Bio-Rad 1610729) at 4°C. Decalcifying solution was refreshed every 24-48 hours until bones were pliable. Bones were processed by the Johns Hopkins Hospital Reference Histology Laboratory and embedded in paraffin blocks. Hematoxylin and eosin staining (H&E) was performed on mounted 5 µm sections of the proximal tibial growth plates. Images were taken with a Nikon 80i microscope under brightfield illumination at 10, 20, and 40x magnification and analyzed using NIS elements software to obtain height measurements of the proliferative zone (PZ), hypertrophic zone (HZ), and growth plate (GP). Measurements were averaged over 3 sites per section, for 4 sections per sample. Cell counts were obtained in the proliferative and hypertrophic zones and averaged for each mouse. Cell size was assessed for hypertrophic chondrocytes using ImageJ and averaged over 15 cells per section, 4 sections per mouse (60 cells total).

### ATDC5 cell culture

ATDC5 cells were sourced from the European Collection of Authenticated Cell Cultures (Sigma-Aldrich 99072806). Cells were propagated at 37°C, 5% CO2, in ATDC5 Complete Media. This consisted of DMEM/F-12 (ThermoFisher Scientific 11320033) supplemented with 5% heat-inactivated fetal bovine serum (ThermoFisher Scientific 16140071), 2 mM L-glutamine (Corning 25-005-CI), 100 U/mL penicillin and 100 μg/mL streptomycin (ThermoFisher Scientific 15140122). Media was changed every 48-72 hours.

### Chondrogenic differentiation

24 hours post-seeding, media was exchanged for ATDC5 Complete Media supplemented with 0.05 mg/mL L-ascorbic acid (Sigma-Aldrich A4403), 10 mM β-glycerophosphoric acid (ThermoFisher Scientific 410991000), and 1x Insulin-Transferrin-Selenium (ITS-G) (ThermoFisher Scientific 41400045). Cells were cultured for up to 28 days at 37°C, 5% CO2, with supplemented media changes every 48-72 hours.

### Generation of *Kdm6a^−/−^* cell lines

The crRNA was designed to target exon 6 of *Mus musculus Kdm6a* with the following guide sequence: 5’-TTCCTTGGCTCGACAAAAGC-3’. Alt-R^®^ CRISPR-Cas9 tracrRNA and Alt-R^®^ custom Cas9 crRNA (Integrated DNA Technologies) were mixed at an equimolar ratio in Nuclease-Free Duplex buffer (Integrated DNA Technologies 11-01-03-01) and allowed to anneal into duplexes. Alt-R^®^ S.p. Cas9 Nuclease V3 (Integrated DNA Technologies 1081058) was introduced for duplex loading. The resulting ribonucleoprotein (RNP) complex was incubated with CRISPRMAX (ThermoFisher Scientific CMAX00001) and Opti-MEM (ThermoFisher Scientific 31985062) for formation of transfection complexes. Meanwhile, ATDC5 cells were rinsed with 1x PBS, dissociated with 0.25% trypsin, 2.21 mM EDTA (Corning 25-053-CL), and collected for counting. Cells were seeded at 2×10^5^ cells per well in 6-well plates concurrently with introduction of the RNP transfection complexes. After 48 hrs at 37°C, 5% CO2, cells were washed and trypsinized. A sample was lysed with QuickExtract™ DNA Extraction Solution (Biosearch Technologies QE09050) to assess editing efficiency with the Alt-R^®^ Genome Editing Detection Kit (Integrated DNA Technologies 1075932). Remaining cells were seeded either in 96-well plates at 1 cell per 50 µL (limiting dilution cloning), or in 10cm dishes at low densities to obtain single colonies. After 10 days of growth, cells were washed with 1x PBS and underwent mild dissociation in warm trypsin for 1 minute. Single colonies were visually identified, isolated, and expanded to establish clonal lines. Successfully edited lines and unedited control lines were identified using the Alt-R^®^ Genome Editing Detection Kit and verified by Sanger sequencing. Forward primer: 5’-AAGATAGAGTGCAGTGGGTTG-3’. Reverse primer: 5’-CAGAAGTCCAAATGCCTTGTAAAT-3’.

### Generation of *Kmt2d^−/−^* cell lines

Please refer to Fahrner et al., 2019 (21).

### Western blot

Cells were washed with cold 1x PBS twice and lysed with RIPA buffer containing protease inhibitor (ThermoFisher Scientific A32953) to isolate total protein. Histones were extracted using a kit following manufacturer’s protocols (Abcam ab113476). The Pierce BCA Protein Assay kit (ThermoFisher Scientific 23225) was used to measure protein concentrations, and 10 μg of each sample was loaded onto a NuPAGE 4 to 12%, Bis-Tris gel (ThermoFisher Scientific NP0336BOX). Following transfer of proteins, PVDF membranes were blocked with 0.5x Intercept (PBS) Blocking Buffer (LI-COR Biosciences 927-70001) for 1 hour at room temperature. Membranes were stained with primary antibodies overnight at 4°C and repeatedly washed with 1x PBS with 0.1% Tween-20 (PBS-T) the following day. Afterwards, membranes were incubated in secondary antibody for one hour at room temperature. Excess antibody was removed with PBS-T washes. Imaging was performed with the LI-COR Odyssey. All antibodies are listed in Supplemental Table 2. ImageJ software was used to quantify blots by densitometry.

### Alcian blue staining

Cells were seeded at a density of 1×10^5^ cells per well in 6-well plates (Corning 353046). Day 7-28 wells underwent chondrogenic differentiation for the specified number of days; Day 0 wells were collected 24 hours post-seeding. Cells were rinsed with 1x PBS and fixed for 5 minutes in 10% neutral buffered formalin (Epredia 9990244). After 3 washes in 1x PBS to remove residual fixative, cells were stained with 1% Alcian blue solution in 3% acetic acid, pH 2.5 (Sigma-Aldrich B8438) for 1 hour in the dark. Unbound dye was removed with 3 washes in 1x PBS. Whole-well images were captured over a transilluminator. An EVOS FL Auto Imaging System (ThermoFisher Scientific) was used to obtain images at 4X, 10X, and 20X magnification. For quantification of the Alcian blue stain, well contents were lysed in a 10% sodium dodecyl sulfate solution (Sigma-Aldrich L3771) for 20 minutes at minimum. Absorbance was measured at 605 nm with a BioTek Synergy 2 Multi-Mode Microplate Reader.

### MTT assay

Cells were seeded at a density of 1×10^3^ cells per well in 96-well plates (Corning 353072). Day 7-28 wells underwent chondrogenic differentiation for the specified number of days; Day 0 wells were collected 24 hours post-seeding. 50 µL of ATDC5 Complete Media and 10 µL of CellTiter 96 AQueous One Solution (Promega G3580) were added to each well. Plates were incubated at 37°C, 5% CO2 for 1.5 hours to allow colorimetric conversion to the formazan product. Absorbance was measured at 490 nm with a BioTek Synergy 2 Multi-Mode Microplate Reader.

### RNA isolation

Cells were seeded at a density of 1×10^5^ cells per well in 6-well plates (Corning 353046) and underwent chondrogenic differentiation for the specified number of days. Samples were collected with TRIzol reagent (ThermoFisher Scientific 15596026) and chloroform. Following phase separation, the aqueous phase containing RNA was put through the RNA Clean & Concentrator-5 kit (Zymo Research R1013) with DNase I digestion to remove genomic DNA contamination.

### RNA-sequencing library preparation

RNA quantity was measured by Qubit RNA Broad Range Assay (ThermoFisher Scientific Q10210). RNA quality was assessed on an Agilent Fragment Analyzer by the Johns Hopkins Single Cell & Transcriptomics Core (JH SCTC) Facility. 1 μg of total RNA per sample was used as input for the NEBNext Ultra II RNA Library Prep kit with Sample Purification Beads (E7775S), in combination with the NEBNext Poly(A) mRNA Magnetic Isolation Module (New England BioLabs E7490L). Libraries were indexed with NEBNext Multiplex Oligos for Illumina (Dual Index Primers Set 1) (New England BioLabs E7600S). Completed libraries underwent quality assessment on the Agilent Fragment Analyzer run by JH SCTC, and were quantified with the NEBNext Library Quant Kit for Illumina (New England BioLabs E7630L). Libraries were pooled to a final concentration of 4 nM and submitted to the Johns Hopkins Genomics Research Core Facility for high-throughput sequencing on the Illumina NovaSeq 6000 platform, using SP flow cells to generate 100-bp paired-end reads.

### Analysis of RNA-sequencing data

The GRCm38 transcriptome was indexed using Salmon 1.9.0 (62), using the GRCm38 genome as the decoy sequence (Mus_musculus.GRCm38.cdna.all.fa.gz and Mus_musculus.GRCm38.dna.primary_assembly.fa.gz, from http://nov2020.archive.ensembl.org/Mus_musculus/Info/Index, release 102). FASTQ files of demultiplexed paired-end reads were mapped using Salmon 1.9.0, with selective alignment and GC bias correction. Transcript quantification files were loaded into R 4.1.2, running Bioconductor 3.14, with tximeta 1.12.4 (63), and transcript-level counts were summarized by gene. Gene-level counts were aggregated for any technical replicates, and filtered for non- or low-expressed genes (median count over samples < 10). Surrogate variable (SV) analysis was performed using sva 3.42.0 (64) for the dual experiment between *Kmt2d^−/−^* and *Kdm6a^−/−^* cell lines, identifying up to 2 SVs in certain contrasts. Differential expression analysis was performed using DESeq2 1.34.0 (65) with a false-discovery rate threshold < 0.1. biomaRt 2.50.3 was used to annotate genes according to Ensembl *Mus musculus* version 102 (66). Differentially expressed genes are listed in Supplemental Appendices 1 and 3-6. For the comparison to prior Kabuki syndrome 1 data, raw reads were downloaded from the Gene Expression Omnibus (GEO), accession number GSE129365, and processed identically. The Mouse Genome Informatics (MGI) database was accessed on July 11, 2023 to download a list of genes falling under the Gene Ontology (GO) category of ‘cartilage development’. Annotations for negative regulators of this process were removed, leaving 200 genes (complete list in Supplemental Appendix 2). A two-sample Wilcoxon test statistic was calculated for the p-values of genes included in this list, and compared against the distribution of test statistics obtained from performing this test on random groups of genes for 10,000 iterations.

### Statistics

With exception of the RNA-seq data analysis (which is described in Methods, ‘Analysis of RNA-sequencing data’), two-tailed, unpaired Student’s t-tests were used with a cutoff of p < 0.05.

### Study approval

The Johns Hopkins Institutional Animal Care and Use Committee approved all animal studies in this project. Procedures were performed following standards described in the NIH *Guide for the Care and Use of Laboratory Animals* (National Academies Press, 2011).

### Data availability

Raw data from RNA-sequencing experiments are in the process of deposition in NCBI’s Gene Expression Omnibus (67). An accession number will be available once assigned. Supplemental Appendices will be posted to BioRxiv. Analysis code will be available on GitHub.

## AUTHOR CONTRIBUTIONS

WYL and JAF isolated long bones from mice; RCR collected and analyzed the micro-CT data. SC imaged growth plate sections and SC and WYL analyzed images. WYL generated *Kdm6a^−/−^* cell lines. CWG performed time courses and characterizations of *Kdm6a^−/−^* cell lines. CWG prepared libraries for RNA-seq. CWG, LB, and KDH analyzed all RNA-seq data. CWG wrote the manuscript. All authors contributed towards editing and approval of the paper. HTB provided *Kdm6a^tm1d/+^* mice. HTB and KDH were involved in helpful discussions. JAF conceived and directed the project and oversaw its completion.

## ACKNOWLEDGMENTS

The *Kdm6a^tm1d/+^* mice were generated and maintained by Giovanni Carosso in the Björnsson lab. The Johns Hopkins Reference Histology Core assisted in processing and sectioning growth plates. The Johns Hopkins Single Cell and Transcriptomics Core Facility (Jake Volk) handled quality control of sequencing libraries. Next-generation sequencing was performed by the Johns Hopkins Genetics Resources Core Facility (David Mohr). BioRender was used to draw figure 1A. This project was primarily funded by K08HD086250 awarded by NIH/NICHD to JAF, a Hartwell Foundation Individual Biomedical Research Award to JAF, a William and Ella Owens Medical Research Foundation grant to JAF, a Baltimore Centre for Musculoskeletal Science 2015 Pilot and Feasibility Award to JAF, and a Johns Hopkins Clinician-Scientist Award to JAF. Additional funding for breeding and maintaining the shared *Kdm6a^tm1d/+^* mice was drawn from NIH DP5OD017877 to HTB, with support from the Louma G. Foundation awarded to HTB. Funding for the RNA-seq experiment comparing KS1 and KS2 was provided by a JHU Bloomberg Distinguished Professorship to Carol W Greider. RCR is supported by Grant BX003724 from the Biomedical Laboratory Research and Development Service of the Veterans Affairs Office of Research and Development. CWG is funded by NIGMS T32GM136577 and a JHU Bloomberg Distinguished Professorship to Carol W Greider.

## Notes

### Competing Interest Statement

HTB is a consultant for Mahzi therapeutics. The other authors have no conflicts of interest.

