## Supplementary material for "Growth retardation in a mouse model of Kabuki syndrome 2 bears mechanistic similarities to Kabuki syndrome 1"

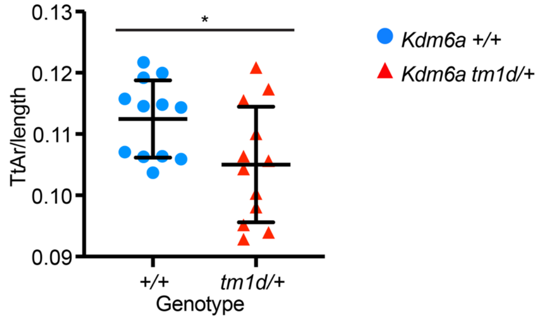


**Supplemental Figure 1: Tissue area of *Kdm6a*$^{tm1d/+}$ and *Kdm6a*$^{+/+}$ littermates, normalized by femur length.** *Kdm6a*$^{tm1d/+}$ mice have decreased tissue area compared to *Kdm6a*$^{+/+}$ even after adjustment for shorter femur lengths. *Kdm6a*$^{+/+}$ n=12; *Kdm6a*$^{tm1d/+}$ n=12. Blue circles: *Kdm6a*$^{+/+}$, red triangles: *Kdm6a*$^{tm1d/+}$. *$p < 0.05$, two-tailed unpaired Student's t-test. All error bars represent mean ± 1 SD. TtAr, total area.

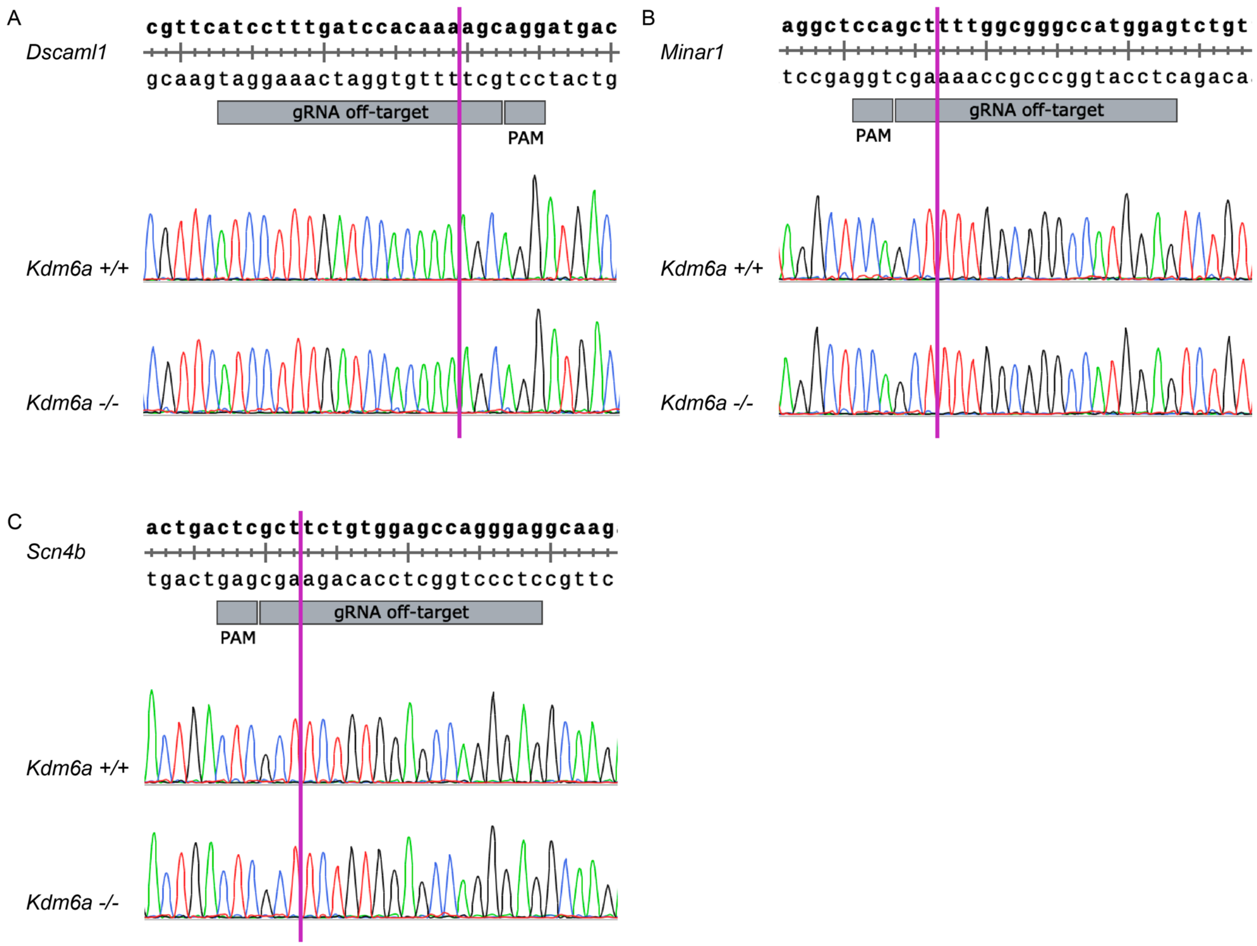


**Supplemental Figure 2: Absence of off-target editing in *Kdm6a*$^{+/+}$ and *Kdm6a*$^{-/-}$ cell lines at top predicted exonic sites.** Representative chromatogram traces from *Kdm6a*$^{+/+}$ and *Kdm6a*$^{-/-}$ lines are shown for: (A) *Dscaml1*, (B) *Minar1*, and (C) *Scn4b*. Purple line indicates predicted off-target Cas9 cleavage site.

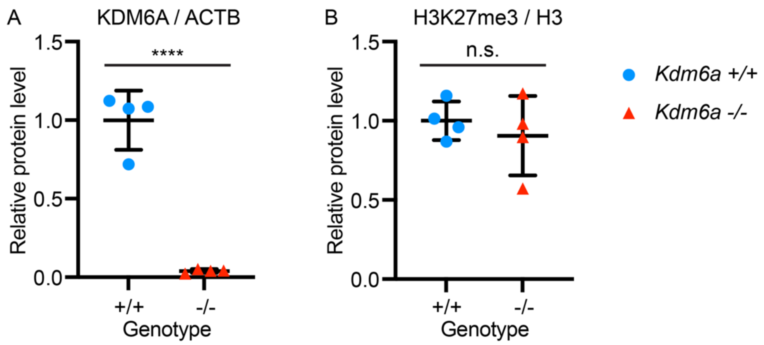


**Supplemental Figure 3: Quantification via densitometry of western blot for Figure 4B.** For both panels, a relative protein level of 1.0 corresponds to the mean densitometry of *Kdm6a*$^{+/+}$ cell lines. (A) *Kdm6a*$^{-/-}$ cells have negligible KDM6A expression, normalized against ACTB loading controls. (B) There is no significant difference in H3K27me3 levels between *Kdm6a*$^{-/-}$ and *Kdm6a*$^{+/+}$ cell lines, with H3 serving as loading control. Blue circles: *Kdm6a*$^{+/+}$, red triangles: *Kdm6a*$^{-/-}$. ****p < 0.0001, two-tailed unpaired Student's t-test. All error bars represent mean ± 1 SD.

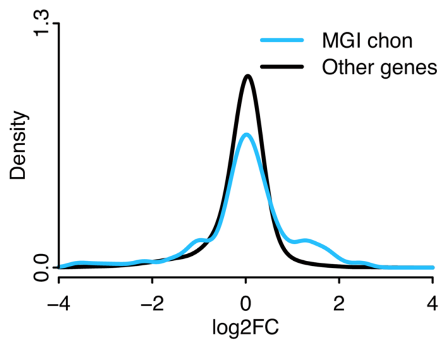


**Supplemental Figure 4: Density distribution of expression fold-changes comparing *Kdm6a*$^{-/-}$ to *Kdm6a*$^{+/+}$ chondrocytes.** A greater proportion of genes annotated by MGI as involved in cartilage development (blue line) exhibit higher absolute fold-changes than non-chondrogenic genes (black line). MGI chon, MGI-annotated chondrogenic genes.

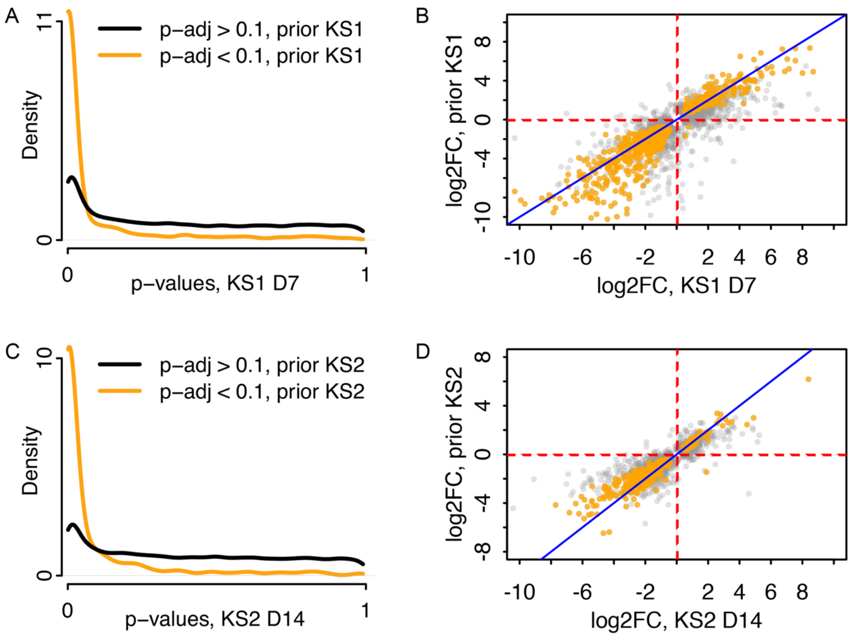


**Supplemental Figure 5: Validation of KS1 and KS2 RNA-seq data by comparison to prior datasets.** (A) Conditional p-value density plot displaying gene-wise p-values from the *Kmt2d*$^{-/-}$ versus *Kmt2d*$^{+/+}$ contrast at Day 7 of chondrogenic differentiation, stratified by significance in a prior dataset comparing *Kmt2d*$^{-/-}$ versus *Kmt2d*$^{+/+}$ cells, also collected at Day 7 (21). Orange: genes that were significant (FDR < 0.1) in the prior dataset; black: non-significant in prior dataset. (B) Scatter plot of gene-wise $\log_2$(fold-changes) in present and prior datasets at Day 7 shows high concordance in fold-change directionalities between datasets. A similar analysis was performed for the *Kdm6a*$^{-/-}$ versus *Kdm6a*$^{+/+}$ contrast at Day 14: (C) conditional p-value histogram and (D) $\log_2$(fold-change) scatter plot.

**Supplemental Table 1: ATDC5 cell line genotypes**

| Line | Genotype | Allele variants |
|---|---|---|
| 10-37 | Compound het | NM_009483.2:c.496del p.(Cys166Valfs*16)<br>NM_009483.2:c.483_493del p.(Asp162Leufs*21) |
| 10-39 | Compound het | NM_009483.2:c.495_496del p.(Phe165Leufs*21)<br>NM_009483.2:c.493_494insA p.(Phe165Tyrfs*22) |
| 96-15 | Compound het | NM_009483.2:c.495_496del p.(Phe165Leufs*21)<br>NM_009483.2:c.496del p.(Cys166Valfs*16) |
| 96-20 | Compound het | NM_009483.2:c.496dup p.(Cys166Leufs*21)<br>NM_009483.2:c.496del p.(Cys166Valfs*16) |
| 96-21 | Compound het | NM_009483.2:c.495_496del p.(Phe165Leufs*21)<br>NM_009483.2:c.496dup p.(Cys166Leufs*21) |
| 10-3 | Wild-type | |
| 10-29 | Wild-type | |
| 96-13 | Wild-type | |
| 96-14 | Wild-type | |
| 96-19 | Wild-type | |

**Supplemental Table 2: Reagents**

| | Species | Manufacturer | Catalog number |
|---|---|---|---|
| anti-H3K27me3 | Rabbit | Millipore Sigma | 07-449 |
| anti-H3 | Rabbit | Cell Signaling Technology | 4499 |
| anti-KDM6A | Rabbit | Cell Signaling Technology | 33510 |
| anti-ACTB | Mouse | Cell Signaling Technology | 3700 |
| IRDye 800CW Donkey anti-Rabbit IgG | Donkey | LI-COR | 926-32213 |
| IRDye 680RD Donkey anti-Mouse IgG | Donkey | LI-COR | 926-68072 |
